# Migrating neurons adapt motility modes to brain microenvironments via a mechanosensor, PIEZO1

**DOI:** 10.1101/2023.01.17.524464

**Authors:** Naotaka Nakazawa, Gianluca Grenci, Yoshitaka Kameo, Noriko Takeda, Tsuyoshi Sawada, Junko Kurisu, Zhejing Zhang, Taiji Adachi, Keiko Nonomura, Mineko Kengaku

**Affiliations:** Institute for Integrated Cell-Material Sciences (KUIAS-iCeMS), Kyoto University, Kyoto, 606-8501, Japan; Mechanobiology Institute, National University of Singapore, Singapore; Biomedical Engineering Department, National University of Singapore, Singapore; Graduate School of Biostudies, Kyoto University, Kyoto, Japan; Institute for Life and Medical Sciences, Kyoto University, Kyoto, Japan; Graduate School of Engineering, Kyoto University, Kyoto, Japan; Department of Life Science and Technology, Tokyo Institute of Technology, Kanagawa, Japan; National Institute for Basic Biology, Aichi, Japan

## Abstract

Migration of newborn neurons is essential for brain morphogenesis and circuit formation, yet controversy exists regarding how neurons generate the driving force against strong mechanical stresses in crowded neural tissues. We found that cerebellar granule neurons adopt differential motility modes in distinct extracellular environments. In 3-dimensional (3D) confinement, actomyosin produces contractile forces at the posterior cell membrane, in addition to the traction force in the leading process that is exclusively observed in 2D cultures. The 3D migration is initiated by activation of a mechanosensitive channel PIEZO1. PIEZO1-induced calcium influx in the soma triggers the PKC-ezrin cascade, which recruits actomyosin to the posterior plasma membrane. Thus, migrating neurons use a mechano-sensing mechanism to activate multiple driving forces to maneuver in irregular brain tissue.

**One-Sentence Summary:** Cerebellar granule neurons use a mechanosensor PIEZO1 to switch migratory modes in confined spaces.

## Main Text

On two-dimensional (2D) substrates, cell migration is driven by the propulsive force of polymerizing cortical actin that is anchored to transmembrane adhesion receptors in the leading edge. However, recent studies have revealed that cells confined in 3D matrices can adopt multiple motility modes based on the physical environment, cell-matrix adhesion, and actomyosin contractility (*1, 2*). The mechanism of 3D migration has been intensively studied using mesenchymal cancer cells, fibroblasts, and immune cells, but is yet to be identified in other migratory cell types, including neural cells.

Postmitotic neurons in the mammalian brain migrate long distances during cortical brain development. In these cells, force generation and transmission also rely on actomyosin dynamics, yet considerable diversity has been found in different models of neuronal migration. For instance, in migrating forebrain interneurons in organotypic slices or Matrigel, non-muscle myosin II (hereafter called myosin) is enriched at the rear of the cell and exerts a pushing force behind the nucleus (*3–6*). In contrast, in migrating cerebellar granule neurons (CGNs) on culture dishes, actomyosin in the leading process generates an adhesion-dependent traction force that pulls the cell soma (*7–10*). The controversy over whether the actomyosin force is generated in the front or rear has been attributed to the differences in neuron-types and assay systems (*11–13*). Here we demonstrate that a migrating neuron is not governed by a single migratory mode but is equipped with multiple mechanisms and can switch between driving forces in 2D and 3D environments.

### Neurons hire differential actomyosin forces in 2D and 3D environments

The radial migration of CGNs in developing brain tissue can be recapitulated in an organotypic cerebellar slice culture (*14, 15*). To ask whether the assay condition influences actomyosin dynamics, we electroporated GFP-tagged myosin regulatory light chain (MRLC-2GFP) in the neonatal mouse cerebellum and observed myosin dynamics in migrating CGNs in the organotypic culture (Fig. S1A). CGNs underwent typical saltatory movements with intermittent forward displacement of the soma. Unlike CGNs in 2D culture, myosin was more broadly distributed and, in many cases, appeared to localize at the rear of the soma prior to a large-amplitude movement (Fig.1, A-C, Fig.S1, B and C).

**Fig. 1.**
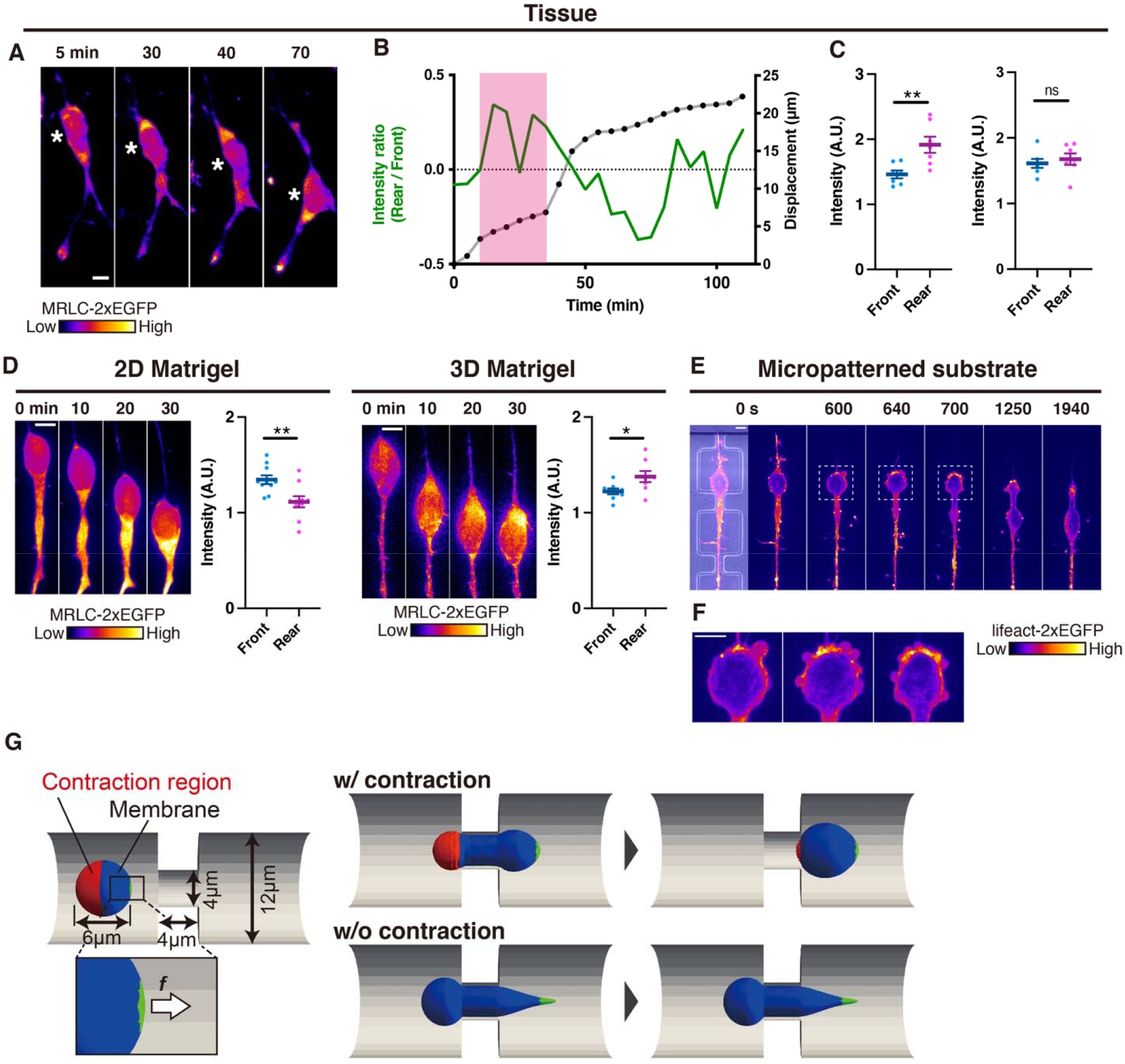
Actomyosin is differentially localized in 2D and 3D microenvironments. (**A**) Image sequence of a CGN migrating in cerebellar tissue isolated from a P10 mouse. (**B**) Position of the soma (approximate oval center indicated by asterisks in A) and relative (rear/front) distribution of myosin (MRLC-GFP) in the neuron shown in A plotted against time. Red shading indicates the period before a large-amplitude movement occurs. (**C**) The rear/front distribution of myosin in migrating CGNs in slice cultures prior to a large-amplitude movement of the soma (left), or averaged over 120 min (right). (**D**) Image sequence and the rear/front comparison of myosin localization in migrating CGNs in 2D (left) and 3D (right) Matrigel culture. (**E**) Micropatterned substrates with repetitive constrictions (left) and an image sequence of a CGN transfected with MRLC-2GFP moving through 3 x 5 μm^2^ constrictions (right 6 images). (**F**) Magnified view of the CGN in (**E**) at 600 s, 640 s, and 700 s. (**G**) Simulation of a deformable spherical object that moves into a narrow tunnel. The red color indicates the area where contraction occurs, and the green color indicates the area where a tensile force is applied. Samples were collected from three independent experiments in (C) and (D). Scale bars, 5 μm.

We next observed the migration of CGNs embedded within 3D Matrigel or plated on its surface (Fig. S1D). In neurons cultured on the Matrigel surface, myosin localized to the cell front and leading process, as was previously observed on 2D substrates (Fig. 1D) (*7, 9, 10*). In contrast, in neurons embedded in Matrigel, myosin appeared more dynamic and frequently enriched at the rear of the cell, as in the cerebellar tissue in slice cultures (Fig. 1D). These results suggest that actomyosin localization in migrating CGNs is differentially regulated in 2D and 3D environments.

We then tested whether such a mode transition is fundamental among neuronal types by examining cortical interneurons. In dissociated interneurons in 3D Matrigel, myosin was highly enriched in the rear of the cell, consistent with previous reports (*3–6*). In contrast, on the Matrigel surface, myosin did not clearly localize in the rear, but instead was more broadly distributed in the neurons (Fig. S1E). Thus, at least two different types of neurons change actomyosin localization depending on the extracellular environment.

To further characterize neuronal migration in a 3D environment, we designed micropatterned channels connected by narrow tunnels that mimic confined extracellular spaces in brain tissues (Fig. S2A). Dissociated CGNs were capable of passing through tunnels with a width of 3 μm (5 μm height), much smaller than the CGN somal diameter of about 6 μm (Fig. 1E, Fig. S2B). Myosin dynamically moved throughout the entire cell soma in neurons situated in a wide channel. When these cells migrated into a narrow tunnel, myosin localized at the rear periphery of the cell until the entire cell soma passed through the tunnel into the next channel (Fig. 1E and Fig. S2C). During the passage of the cell soma through the tunnel, we observed that transient membrane blebs actively formed at the actin-rich cell rear, implying actomyosin contractility at the posterior plasma membrane (*16*)(Fig. 1F). These results suggest that neurons localize actomyosin at the rear plasma membrane to exert a contractile force assisting the passage through a narrow constriction.

We next performed a computational simulation to verify whether actomyosin contractility in the cell rear is effective in confined migration. We assumed a cell resembles a deformable and slightly compressible spherical object covered with a hyperelastic membrane. The model cell entered a narrow tunnel by tensile force on the front membrane with or without the contraction at the rear edge (Supplementary Text and Table S1). With only the front force, the cell deformed and moved forward until the front edge entered the constriction, but the rear half stopped at the constriction. In contrast, when the contraction of the membrane at the rear edge was combined with the front force, the entire cell successfully entered and passed through the narrow tunnel (Fig. 1G). Collectively, the simulation results suggest that actomyosin contraction at the cell rear can produce forward propulsion that enables neurons to pass through narrow constrictions.

### PIEZO1 signaling triggers actomyosin force transmission in 3D confinement

In order to identify the mechanism regulating the posterior localization and contraction of actomyosin in a confined microenvironment, we performed a pharmacological screen in a transwell assay where cells migrated through polycarbonate membranes with different pore sizes (Fig. 2A). The majority of CGNs seeded on the membranes underwent transwell migration through both confined 3-μm pores (67% at 6 hr) and permissive 8-μm pores (85% at 6 hr)(Fig. S3, A and B). The inhibition of actin turnover (cytochalasin D, latrunculin B, or jasplakinolide) or myosin activity (blebbistatin) strongly reduced transmigration through both 3-μm and 8-μm pores, supporting the notion that actomyosin dynamics is indispensable for neuronal migration in both confined and non-confined spaces (Fig. 2B). Among a set of molecules tested under confined and non-confined conditions, we found that treatment with GsMTx4, a peptide inhibitor of mechanosensitive ion channels (MSCs), specifically reduced transmigration of CGNs through 3-μm pores, but not 8-μm pores (Fig. 2C) (*17, 18*). As GsMTx4 inhibits both Transient Receptor Potential (TRP) and PIEZO families, we asked which class of MSCs was responsible for confined migration. Inhibition of TRPC channels by SKF96365 or the TRPV4 channel by HC-067047 had no overt differential effects on migration through 3-μm and 8-μm pores (Fig. 2D)(*19, 20*). In contrast, conditional deletion of PIEZO1 in CGNs (NeuroD1-Cre; Piezo1^flox/flox^, hereafter called PIEZO1 cKO) strongly reduced transwell migration through 3-μm pores, but had little effect on migration through 8-μm pores (Fig. 2E) (*21*). These results suggest that mechanical stress during penetration into the narrow pores activates PIEZO1 and evokes the molecular signal necessary for confined migration.

**Fig. 2.**
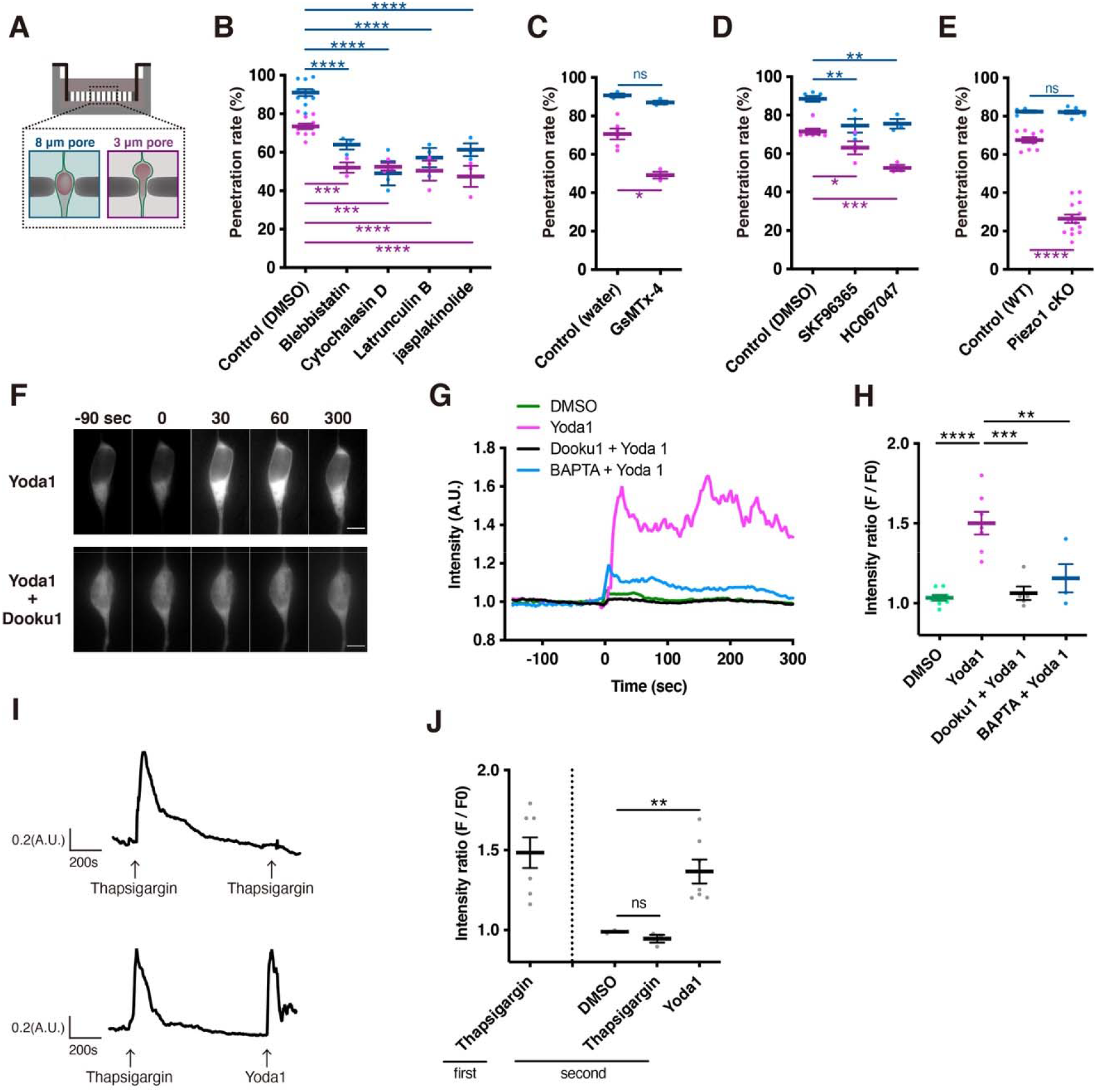
PIEZO1 is active in migrating CGNs. (**A**) Transwell assay. (**B**) Treatment with inhibitors of actomyosin dynamics. The percentages of cells that migrated to the bottom side through 3-μm (purple) or 8-μm (blue) pores (penetration rates) were calculated. (**C-E**) The effects of a mechanosensitive channel blocker GsMTx4 (**C**), TRP channel inhibitors (**D**) and conditional deletion of PIEZO1 (**E**). (**F**) Ca^2+^ elevation in cultured CGNs treated with Yoda1(100 μM) with or without its antagonist Dooku1 (100 μM). Drugs were added at time 0. Scale bars, 5 μm. (**G**) The GCAMP6s ΔF/F transients in the soma upon treatment with Yoda1 in the presence or absence of Dooku1 or BAPTA. (**H**) Mean ratio of peak amplitudes of GCAMP6s ΔF/F transients. (**I**) Ca^2+^ elevation evoked by 2 treatments with thapsigargin (top) and a treatment with thapsigargin followed by Yoda1 (bottom). (**J**) Mean ratio of peak amplitudes of GCAMP6s ΔF/F transients. Samples were collected from three independent experiments.

*PIEZO1 mRNA* expression in developing CGNs was confirmed by RNAscope *in situ* hybridization and RNA-seq analysis (Fig. S3, C and D and Data S1). We examined PIEZO1 activity in CGNs by applying a specific agonist, Yoda1, in a dissociated culture (*22*). In the presence of external Ca^2+^ (1.8 mM), Yoda1 treatment induced a slow and prolonged increase in the intracellular Ca^2+^ concentration ([Ca^2+^]_i_) as visualized by a GCaMP6s probe (*23*)(Fig. 2, F and G). The increase in [Ca^2+^]_i_ by Yoda1 treatment was inhibited by the selective PIEZO1 inhibitor Dooku1 or an extracellular Ca^2+^ chelator BAPTA, indicating that the activated PIEZO1 allows Ca^2+^ entry into the CGN cytoplasm (Fig. 2, F-H). Previous studies have shown that PIEZO1 localizes to both the plasma membrane and the endoplasmic reticulum (ER) and mediates Ca^2+^ release from the ER as well as the influx of extracellular Ca^2+^ (*24, 25*). To identify the source of the [Ca^2+^]_i_ increase induced by Yoda1, we pretreated the CGNs with an ER Ca^2+^-ATPase blocker thapsigargin to exhaust the ER Ca^2+^ store before the addition of Yoda1. Thapsigargin pretreatment induced a transient [Ca^2+^]_i_ rise, while the second thapsigargin administration at 15 min caused little or no response in the same cell, confirming that the pretreatment depleted Ca^2+^ from the ER (Fig. 2I, top). In contrast, Yoda1 treatment at 15 min after the thapsigargin pretreatment induced a large [Ca^2+^]_i_ elevation, indicating that PIEZO1 induces the bulk of the Ca^2+^ influx from the extracellular environment (Fig. 2, I, bottom, and J).

Notably, the Yoda1-induced Ca^2+^ elevation was followed by the surface accumulation of actomyosin and twitching of the cell soma, suggesting that the activation of PIEZO1 induced actomyosin translocation to the plasma membrane (Fig. 3A). We therefore sought to identify the signaling pathway downstream of PIEZO1 using transwell assays. We first confirmed that BAPTA or EGTA treatment specifically attenuated migration through 3-μm pores but not 8-μm pores (Fig. 3B). We found that pharmacological inhibition of Ca^2+^-dependent protein kinase C (PKC) by Gö6983 (PKCα, β, γ) or LY333531 (PKCβ), or ezrin by NSC668394, also produced differential effects on migration through 3-μm and 8-μm pores (Fig. 3C). Furthermore, phosphorylation of ezrin at threonine 567 (T567) was significantly elevated in Yoda1-treated CGNs compared to the untreated control (Fig. 3D). The phosphorylation at T567 is known to be regulated by Ca^2+^-dependent PKCs (*26–28*). Indeed, elevation of ezrin T567 phosphorylation by Yoda1 treatment was inhibited when the cells were preincubated with PKCβ inhibitor LY333531. Furthermore, the basal level of ezrin phosphorylation was downregulated in PIEZO1 cKO CGNs (Fig. 3D). These results suggest that PIEZO1-induced Ca^2+^ influx activates Ca^2+^-dependent PKCβ, which in turn phosphorylates ezrin in CGNs.

**Fig. 3.**
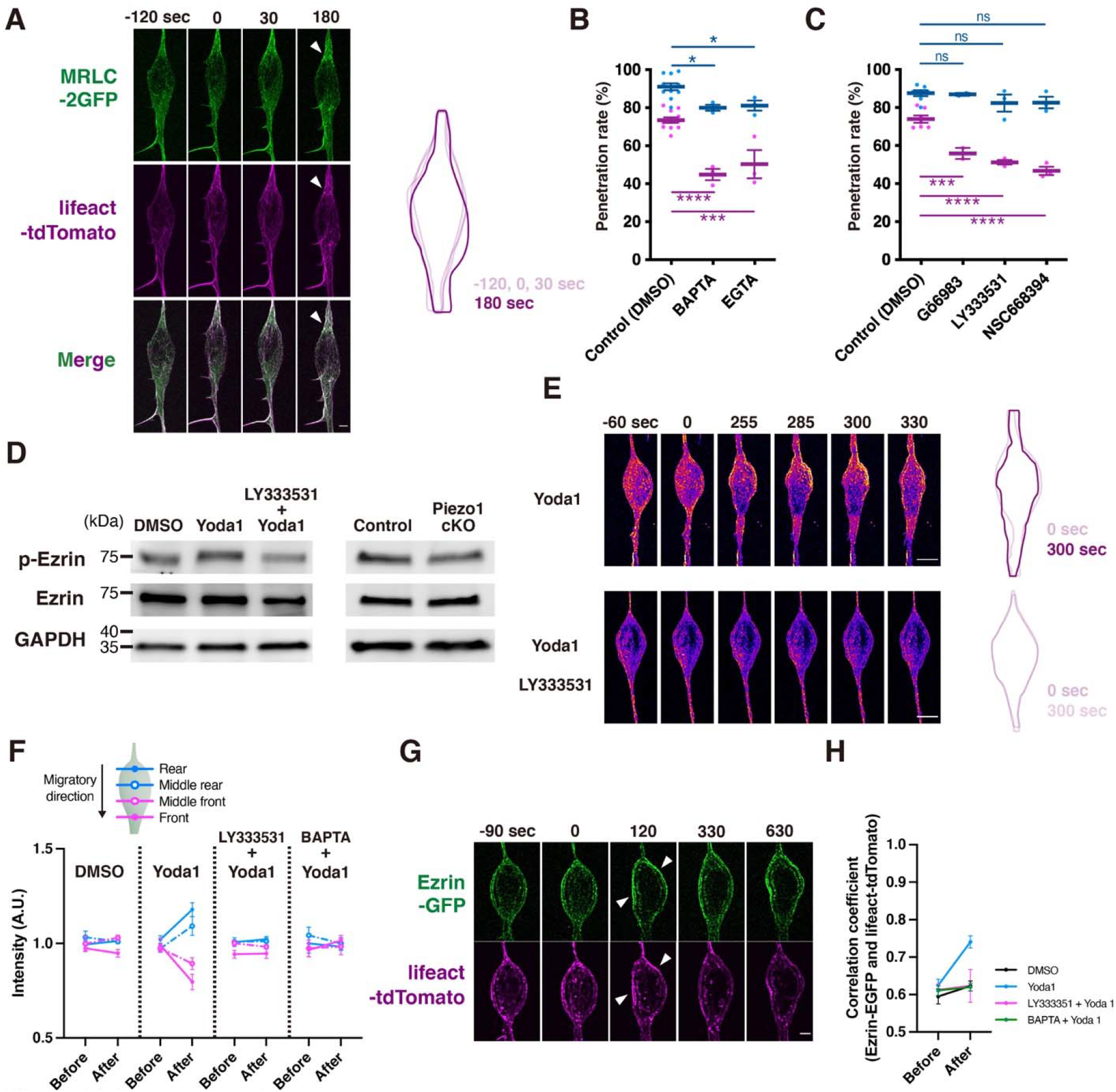
PIEZO1 induces plasma membrane recruitment of actomyosin via the PKC-ezrin pathway. (**A**) Time-lapse sequence of MRLC-GFP and lifeact-tdTomato signals in CGNs upon Yoda1 treatment (left). Colored outlines mark the periphery of the soma at various time points (right). (**B, C**) Percentages of cells that passed through 3- or 8-μm transwell pores in the presence of Ca^2+^chelators (**B**), and inhibitors of Ca^2+^-dependent PKCs or ezrin (**C**). (**D**) Phosphorylation and activation of ezrin in CGNs treated with Yoda1 with or without LY333531 (left) and in CGNs from wildtype and PIEZO1 cKO mice (right). Western blotting was performed with antibodies against phosphoT576 ezrin (p-Ezrin), total ezrin (Ezrin) and GAPDH. (**E**) Time-lapse sequence of ezrin-GFP signals in CGNs upon treatment with Yoda1 (left). Cells were pretreated with or without LY333531. Traces at right are shape changes of the soma after Yoda1 treatment. (**F**) Translocation of ezrin to the plasma membrane after drug treatments. The cell soma is subdivided into 4 anteroposterior parts. (**G**) Time-lapse sequence of ezrin-GFP (top) and lifeact-tdTomato (bottom) in CGNs upon treatment with Yoda1. (**H**) Colocalization of ezrin and actomyosin. Samples were collected from three independent experiments. Scale bars, A, G, 2 μm; B, 5 μm.

The ERM (ezrin-radixin-moesin) proteins translocate to the plasma membrane upon phosphorylation and tether the actin cytoskeleton to the plasma membrane (*29, 30*). We hypothesized that Ca^2+^ signaling induced by PIEZO1 leads to the activation and translocation of ezrin and actomyosin to the plasma membrane and induces the somal contraction. We observed that Yoda1 treatment induced significant translocation of ezrin-EGFP to the plasma membrane at the cell rear, followed by somal twitching (Fig. 3, E and F). Strikingly, actin filaments visualized by lifeact-tdTomato were recruited at the sites of ezrin-EGFP localization upon Yoda1 treatment (Fig. 3G and H). Pretreatment with BAPTA or LY333531 abolished not only the translocation of ezrin and actin but cell soma twitching in response to Yoda1 treatment (Fig. 3, E-H, Fig. S4). These data strongly suggest that PIEZO1-mediated Ca^2+^ influx facilitates actomyosin force transmission at the posterior cell membrane through activation of the PKCβ-ezrin pathway.

### PIEZO1 activity is required for neuronal migration in confined brain tissue

We further examined the functional significance of PIEZO1 in CGN migration in a confined microenvironment. The migratory capacity of CGNs from wildtype and PIEZO1 cKO mice was analyzed using a reaggregate culture (*15*) either inside (3D) or on the surface (2D) of Matrigel. There was no apparent difference in migratory capacity between wildtype and PIEZO1 cKO neurons in the 2D culture on the Matrigel surface, whereas PIEZO1 cKO neurons exhibited a significant delay in 3D migration within Matrigel (Fig. 4A). We next transfected dissociated wildtype CGNs with either GFP-ezrin or MRLC-2GFP and subjected them to 2D and 3D Matrigel cultures. Ezrin and myosin were localized at the periphery of the posterior part of the soma during migration in the 3D matrix, while no pronounced localization was seen in cells migrating in the 2D environment (Fig. 4, B and C). In contrast, neither ezrin nor myosin localized posteriorly in PIEZO1-deficient cells in 3D culture, supporting the notion that PIEZO1 is activated in confined spaces and induces ezrin and myosin translocation to the plasma membrane at the cell rear (Fig. 4, B and C).

**Fig. 4.**
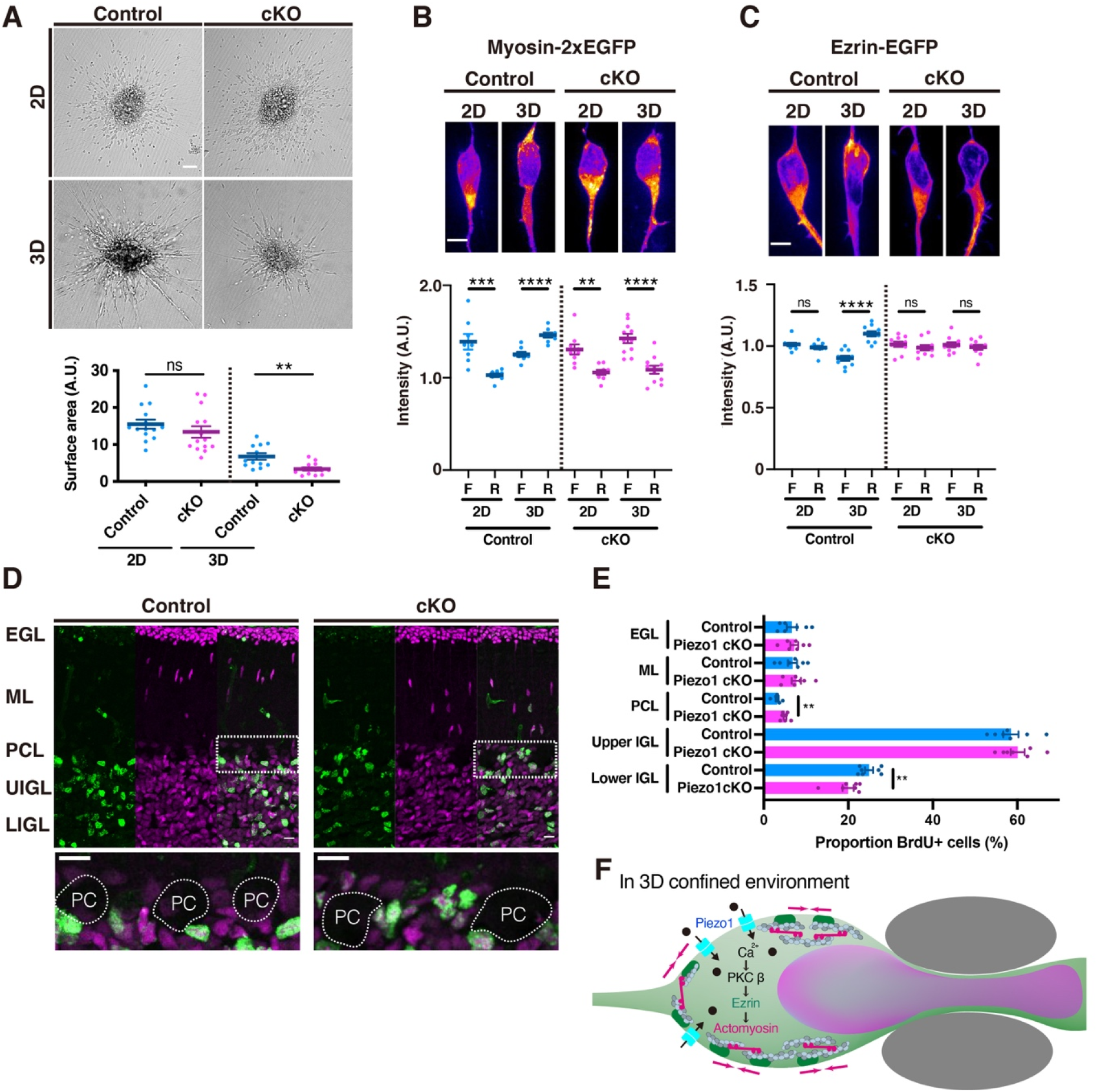
PIEZO1 signaling is required for CGN migration in confined microenvironments. (**A**) Migration of CGNs from wildtype and PIEZO1 cKO mice in reaggregate cultures. Reaggregates were plated on the surface (2D) or embedded in Matrigel (3D). Bottom graphs show migration distance of CGNs from the reaggregate periphery in each condition. (**B, C**) Snapshot images (top) and the rear/front distribution (bottom) of MRLC-2GFP (**B**) and ezrin-GFP (**C**) in migrating CGNs from wildtype mice and PIEZO1 cKO in 2D and 3D Matrigel culture. (**D**) BrdU(green) and Pax6 (magenta) labeling of P13 cerebella after BrdU injection at P9. A magnified view of the PC layer is shown in bottom panels. (**E**) Proportion of BrdU+ CGNs in each layer. (**F**) Signaling cascade of 3D migration of CGNs. Samples were collected from three independent experiments. Scale bars: A, 50 μm; B,C,D, 10 μm.

We then asked whether PIEZO1 function is involved in CGN migration in vivo. CGNs are born in the external granule layer (EGL) and undergo radial migration toward the internal granule layer (IGL) during the first to third postnatal weeks. There was no apparent difference in the gross morphology of the cerebellar cortices in wildtype and PIEZO1 cKO mice (Fig. S5). We administrated BrdU to label granule cell progenitors at P9 and tracked BrdU-positive CGNs in P13 cerebelli from wildtype and PIEZO1 cKO mice. We found a slight but significant increase in the number of CGNs that had stopped at the Purkinje cell layer (PCL) in PIEZO1 cKO mice. In contrast, CGNs that reached the lower IGL were fewer in PIEZO1 cKO mice, indicating that CGN migration is delayed in crowded tissue space in the absence of PIEZO1 (Fig. 4, D and E).

## Discussion

In summary, we have demonstrated that neurons switch their migration modes by sensing the extracellular microenvironment in the developing brain. Migration is driven by the adhesion-based traction generated in the cell front in a non-confined environment, but when CGNs encounter a constricted space, they exert an actomyosin contractile force at the cell rear to squeeze into the constriction. The multi-step processes of confined migration of neurons depicted by the present study include: (1) the entry of the cell front into a constriction; (2) increase in internal pressure and membrane tension in the cell soma; (3) PIEZO1 activation and Ca^2+^ influx; (4) activation of Ca^2+^-dependent PKC; (5) ezrin-actomyosin coupling to the posterior plasma membrane; (6) forward propulsion by the actomyosin contractile force at the rear (Fig. 4F). Piezo1 has been implicated in actin remodeling during cell migration, but its role in the rear-biased actomyosin contraction in 3D migration has not been reported (*16, 31, 32*). Notably, ezrin translocation was strongly biased to the posterior membrane by bath-applied Yoda1 in non-confined spaces, suggesting that asymmetric shape and/or molecular distribution in the polarized neuron also contribute to local activation of ezrin and actomyosin at the cell rear (*2*). We observe that cortical interneurons also switch between multiple migration modes, yet whether they use the same signaling pathway as CGNs is unknown. It has been shown that ERM phosphorylation is regulated by another MSC TRPV4 in migrating breast cancer cells (*33*). TRPV4 activity has also been implicated in increasing osmotic pressure in the cell front to propel forward migration of mesenchymal stem cells in a confined space (*34*). Thus, strategies for 3D migration might be more diverse among various cell types than currently understood.

## Acknowledgments

We thank F. Margadant, and H. Xian for invaluable discussion, G. Zimmer-Bensch, D. Pensold, M. Sawada, and K. Sawamoto for culture ptotocols, F. Ishidate, H. Umeshima, and iCeMS Analysis Center for technical assistance, A. Patapoutian for kind gift of Piezo^flox/flox^ mouse line.

## Author contributions

NN and MK conceived the project and designed the experiments. GG designed and generated micropatterned substrates. NN, JK, NT and KN performed the experiments. YK, TS and TA performed modeling. MK and NN wrote the manuscript with input from all authors.

## Data and materials availability

All data are available in the main text or the supplementary materials.

## Supplementary Materials

Materials and Methods

Supplementary Text

Figs. S1 to S5

Table S1

References (*1–12*)

Data S1

## References

1. K. M. Yamada, M. Sixt, Mechanisms of 3D cell migration. Nat. Rev. Mol. Cell Biol. 20, 738–752 (2019).

2. A. C. Callan-Jones, R. Voituriez, Actin flows in cell migration: from locomotion and polarity to trajectories. Curr. Opin. Cell Biol. 38, 12–17 (2016).

3. B. T. Schaar, S. K. McConnell, Cytoskeletal coordination during neuronal migration. Proc. Natl. Acad. Sci. U. S. A. 102, 13652–13657 (2005).

4. A. Bellion, J.-P. Baudoin, C. Alvarez, M. Bornens, C. Métin, Nucleokinesis in tangentially migrating neurons comprises two alternating phases: forward migration of the Golgi/centrosome associated with centrosome splitting and myosin contraction at the rear. J. Neurosci. 25, 5691–5699 (2005).

5. F. J. Martini, M. Valdeolmillos, Actomyosin contraction at the cell rear drives nuclear translocation in migrating cortical interneurons. J. Neurosci. 30, 8660–8670 (2010).

6. R. Shinohara, D. Thumkeo, H. Kamijo, N. Kaneko, K. Sawamoto, K. Watanabe, H. Takebayashi, H. Kiyonari, T. Ishizaki, T. Furuyashiki, S. Narumiya, A role for mDia, a Rho-regulated actin nucleator, in tangential migration of interneuron precursors. Nat. Neurosci. 15, 373–80, S1-2 (2012).

7. J. Jiang, Z.-H. Zhang, X.-B. Yuan, M.-M. Poo, Spatiotemporal dynamics of traction forces show three contraction centers in migratory neurons. J. Cell Biol. 209, 759–774 (2015).

8. D. J. Solecki, N. Trivedi, E.-E. Govek, R. A. Kerekes, S. S. Gleason, M. E. Hatten, Myosin II motors and F-actin dynamics drive the coordinated movement of the centrosome and soma during CNS glial-guided neuronal migration. Neuron. 63, 63–80 (2009).

9. N. Trivedi, D. R. Stabley, B. Cain, D. Howell, C. Laumonnerie, J. S. Ramahi, J. Temirov, R. A. Kerekes, P. R. Gordon-Weeks, D. J. Solecki, Drebrin-mediated microtubule-actomyosin coupling steers cerebellar granule neuron nucleokinesis and migration pathway selection. Nat. Commun. 8, 14484 (2017).

10. H. Umeshima, K.-I. Nomura, S. Yoshikawa, M. Hörning, M. Tanaka, S. Sakuma, F. Arai, M. Kaneko, M. Kengaku, Local traction force in the proximal leading process triggers nuclear translocation during neuronal migration. Neurosci. Res. 142, 38–48 (2019).

11. T. Minegishi, N. Inagaki, Forces to Drive Neuronal Migration Steps. Frontiers in Cell and Developmental Biology. 8, 863 (2020).

12. M. Kengaku, Cytoskeletal control of nuclear migration in neurons and non-neuronal cells. Proc. Jpn. Acad. Ser. B Phys. Biol. Sci. 94, 337–349 (2018).

13. N. Trivedi, D. J. Solecki, Neuronal migration illuminated: a look under the hood of the living neuron. Cell Adh. Migr. 5, 42–47 (2011).

14. Y. K. Wu, H. Umeshima, J. Kurisu, M. Kengaku, Nesprins and opposing microtubule motors generate a point force that drives directional nuclear motion in migrating neurons. Development. 145 (2018), doi:10.1242/dev.158782.

15. H. Umeshima, T. Hirano, M. Kengaku, Microtubule-based nuclear movement occurs independently of centrosome positioning in migrating neurons. Proc. Natl. Acad. Sci. U. S. A. 104, 16182–16187 (2007).

16. R. Poincloux, O. Collin, F. Lizárraga, M. Romao, M. Debray, M. Piel, P. Chavrier, Contractility of the cell rear drives invasion of breast tumor cells in 3D Matrigel. Proc. Natl. Acad. Sci. U. S. A. 108, 1943–1948 (2011).

17. S.-Y. Lee, R. MacKinnon, A membrane-access mechanism of ion channel inhibition by voltage sensor toxins from spider venom. Nature. 430, 232–235 (2004).

18. T. M. Suchyna, J. H. Johnson, K. Hamer, J. F. Leykam, D. A. Gage, H. F. Clemo, C. M. Baumgarten, F. Sachs, Identification of a peptide toxin from Grammostola spatulata spider venom that blocks cation-selective stretch-activated channels. J. Gen. Physiol. 115, 583–598 (2000).

19. J. E. Merritt, W. P. Armstrong, C. D. Benham, T. J. Hallam, R. Jacob, A. Jaxa-Chamiec, B. K. Leigh, S. A. McCarthy, K. E. Moores, T. J. Rink, SK&F 96365, a novel inhibitor of receptor-mediated calcium entry. Biochem. J. 271, 515–522 (1990).

20. W. Everaerts, X. Zhen, D. Ghosh, J. Vriens, T. Gevaert, J. P. Gilbert, N. J. Hayward, C. R. McNamara, F. Xue, M. M. Moran, T. Strassmaier, E. Uykal, G. Owsianik, R. Vennekens, D. De Ridder, B. Nilius, C. M. Fanger, T. Voets, Inhibition of the cation channel TRPV4 improves bladder function in mice and rats with cyclophosphamide-induced cystitis. Proc. Natl. Acad. Sci. U. S. A. 107, 19084–19089 (2010).

21. S. M. Cahalan, V. Lukacs, S. S. Ranade, S. Chien, M. Bandell, A. Patapoutian, Piezo1 links mechanical forces to red blood cell volume. Elife. 4 (2015), doi:10.7554/eLife.07370.

22. R. Syeda, J. Xu, A. E. Dubin, B. Coste, J. Mathur, T. Huynh, J. Matzen, J. Lao, D. C. Tully, I. H. Engels, H. M. Petrassi, A. M. Schumacher, M. Montal, M. Bandell, A. Patapoutian, Chemical activation of the mechanotransduction channel Piezo1. Elife. 4 (2015), doi:10.7554/eLife.07369.

23. T.-W. Chen, T. J. Wardill, Y. Sun, S. R. Pulver, S. L. Renninger, A. Baohan, E. R. Schreiter, R. A. Kerr, M. B. Orger, V. Jayaraman, L. L. Looger, K. Svoboda, D. S. Kim, Ultrasensitive fluorescent proteins for imaging neuronal activity. Nature. 499, 295–300 (2013).

24. B. Coste, J. Mathur, M. Schmidt, T. J. Earley, S. Ranade, M. J. Petrus, A. E. Dubin, A. Patapoutian, Piezo1 and Piezo2 are essential components of distinct mechanically activated cation channels. Science. 330, 55–60 (2010).

25. S. A. Gudipaty, J. Lindblom, P. D. Loftus, M. J. Redd, K. Edes, C. F. Davey, V. Krishnegowda, J. Rosenblatt, Mechanical stretch triggers rapid epithelial cell division through Piezo1. Nature. 543, 118–121 (2017).

26. T. Ng, M. Parsons, W. E. Hughes, J. Monypenny, D. Zicha, A. Gautreau, M. Arpin, S. Gschmeissner, P. J. Verveer, P. I. H. Bastiaens, P. J. Parker, Ezrin is a downstream effector of trafficking PKC–integrin complexes involved in the control of cell motility. EMBO J. 20, 2723–2741 (2001).

27. J. Clucas, F. Valderrama, ERM proteins in cancer progression. J. Cell Sci. 128, 1253 (2015).

28. K. Endo, S. Kondo, J. Shackleford, T. Horikawa, N. Kitagawa, T. Yoshizaki, M. Furukawa, Y. Zen, J. S. Pagano, Phosphorylated ezrin is associated with EBV latent membrane protein 1 in nasopharyngeal carcinoma and induces cell migration. Oncogene. 28, 1725–1735 (2009).

29. O. Turunen, T. Wahlström, A. Vaheri, Ezrin has a COOH-terminal actin-binding site that is conserved in the ezrin protein family. J. Cell Biol. 126, 1445–1453 (1994).

30. R. G. Fehon, A. I. McClatchey, A. Bretscher, Organizing the cell cortex: the role of ERM proteins. Nat. Rev. Mol. Cell Biol. 11, 276–287 (2010).

31. B. Canales Coutiño, R. Mayor, The mechanosensitive channel Piezo1 cooperates with semaphorins to control neural crest migration. Development. 148 (2021), doi:10.1242/dev.200001.

32. Y.-J. Liu, M. Le Berre, F. Lautenschlaeger, P. Maiuri, A. Callan-Jones, M. Heuzé, T. Takaki, R. Voituriez, M. Piel, Confinement and low adhesion induce fast amoeboid migration of slow mesenchymal cells. Cell. 160, 659–672 (2015).

33. W. H. Lee, L. Y. Choong, N. N. Mon, S. Lu, Q. Lin, B. Pang, B. Yan, V. S. R. Krishna, H. Singh, T. Z. Tan, J. P. Thiery, C. T. Lim, P. B. O. Tan, M. Johansson, C. Harteneck, Y. P. Lim, TRPV4 Regulates Breast Cancer Cell Extravasation, Stiffness and Actin Cortex. Sci. Rep. 6, 27903 (2016).

34. H.-P. Lee, F. Alisafaei, K. Adebawale, J. Chang, V. B. Shenoy, O. Chaudhuri, The nuclear piston activates mechanosensitive ion channels to generate cell migration paths in confining microenvironments. Sci Adv. 7 (2021), doi:10.1126/sciadv.abd4058.

