## Supplementary Materials for "Migrating neurons adapt motility modes to brain microenvironments via a mechanosensor, PIEZO1"

### Newborn neurons adapt migratory mode to brain microenvironments via a mechanosensor, PIEZO1

**This PDF file includes:**

Materials and Methods
Supplementary Text
Figs. S1 to S5
Tables S1

**Other Supplementary Materials for this manuscript include the following:**

Data S1

### Materials and Methods

#### Mice

Mice were handled in accordance with the guidelines of the Animal Experiment Committee of Kyoto University. Timed-pregnant ICR females were purchased from Japan SLC. *PIEZO1*$^{flox/flox}$ mice were generated as previously described (*1*). NeuroD1-Cre mice were obtained from MMRRC at UC Davis [StockTg (Neurod1-cre)RZ24Gsat/Mmucd] and backcrossed to the C57BL/6 strain. Postmitotic neuron-specific PIEZO1 mutant mice were obtained by crossing *PIEZO1*$^{+/flox}$ mice with NeuroD1-Cre$^{+/Cre}$ mice, and littermates derived from *PIEZO1*$^{flox/flox}$ and *PIEZO1*$^{flox/flox}$; NeuroD1-Cre$^{+/Cre}$ mice on the pure C57BL/6 genetic background were used for subsequent studies. Newborn homozygous floxed PIEZO1 littermates were genotyped for presence or lack of NeuroD1-Cre either on postnatal day 4-5 for use in primary cultures or immunohistochemistry experiments. No distinction between male and female pups were made for experiments. Mice were kept in a 12 hr dark/light cycle at 23±3ºC/50% humidity in standard SPF housing.

#### Reagents

Chemicals and the working concentrations used in the pharmacological assays are as follows: blebbistatin (Sigma, B0560, 50 µM), cytochalasin D (TOCRIS, 1233, 10 µM), latrunculin B (KOM, AG-CN2-0031-M001, 1 µM), jasplakinolide (CALBIOCHEM, 420107, 5 µM), GsMTx4 (Abcam, ab141871, 10 µM), SKF96365 (CALBIOCHEM, 567310, 100 µM), HC-067047 (Millipore, 616521, 1 µM), Yoda1 (Sigma, SML1558, 100 µM), Dooku1 (TOCRIS, 6568, 100 µM), BAPTA, extracellular calcium chelator (Abcam, ab144924, 5 µM), EGTA (ThermoFisher, E1219, 100 µM), thapsigargin (Abcam, ab120286, 2 µM), Gö6983 (CALBIOCHEM, 365251, 1 µM), LY333531 (Enzo, ALX-270-348, 10 µM), and NSC668394 (Millipore, 341216, 10 µM).

#### Genotyping

Tail clippings were digested in 50 mM NaOH for 10 minutes at 95ºC. The *PIEZO1*$^{flox}$ allele (LoxP-frt check) was identified using primers 5'CTTGACCTGTCCCCTTCCCCATCAAG3', 5'AGGTTGCAGGGTGGCATGGCTCTTTTT3' and 5'CAGTCACTGCTCTTAACCATTGAGCCATCTC3', and the NeuroD1-Cre$^{cre}$ allele using primers 5' TAGGATTAGGGAGAGGGAGCTGAA 3' and 5' CGGCAAACGGACAGAAGCATT 3', respectively. PCR was performed using KOD FX polymerase kit (Toyobo).

#### In vivo electroporation and image acquisition

P7 or P8 mice were anesthetized on ice and injected with plasmid DNAs diluted in Tris-EDTA buffer with a 33-gauge needle connected to the anode. Tweezer-type electrodes connected to the cathode were placed on the occipital regions. Six electric pulses of 70 mV for 50 ms duration were applied with 150 ms intervals using CUY21 (Nepagene). The pups were revived at 37°C and returned to the litter. Two days after electroporation, cerebella were removed and embedded in 3.5% agarose and sectioned into 300 µm-thick coronal slices with a vibratome. Slices placed on Millicell-CM (Millipore) were mounted in collagen gel and soaked in the media (60% BME, 25% Earle's balanced salt, 15% horse serum, 3 mM L-glutamine, 1 mM sodium pyruvate, 5.6 g/L glucose, 1.8 g/L sodium bicarbonate, 1x N-2 supplement). The tissue was kept in an incubator chamber attached to an upright microscope stage BX61WI (Olympus) at 37°C with 85% $O_2$ / 5% $CO_2$ flow. Images were obtained with FV1000 equipped with a confocal GaAsP detector every 5 min through a 60x water-immersion objective (N.A. 1.1).

Matrigel culture

For primary CGC cultures, cerebella from postnatal day 4-6 mouse littermates of both sexes were dissected in HBSS (Gibco), pooled together, and dissociated using the Neuron Dissociation Kit (Wako Pure Chemical Industries, Ltd). Dissociated neurons were transfected with 6 µg of plasmid DNAs by using lipofectamine 2000 (Thermo Fisher Scientific) according to the manufacturer's instruction. Transfected cells were incubated in non-coated dishes for 8 hours to make cell aggregates. For 3D cultures, resultant aggregates were suspended in 60% Matrigel (Growth Factor Reduced, Corning) in the medium (BME with 26.4 mM D-glucose, 25 mM sodium bicarbonate, 1% bovine serum albumin, 1x N-2 supplement and Penicillin-Streptomicin) and plated in glass-bottom 35-mm culture plates (Iwaki) and then incubated at 37°C for 30 min to allow a gel to form. Each well was then flooded with the culture medium. For 2D cultures, 60% Matrigel was plated into glass-bottom plates and solidified at 37°C for 30 min. Reaggregates were suspended in the culture medium and plated on top of the gel layer. Neurons were incubated in 37°C/5% $CO_2$ for 24h and subjected for live-cell imaging.

Interneurons were prepared from the medial ganglionic eminence (MGE) of embryonic day 14.5 mice and dissociated in 0.04% trypsine in HBSS/0.65% Glucose as described in (*2*). Dissociated interneurons were subjected for transfection and Matrigel culture as described above.

Transwell assay

Dissociated CGCs were transferred in culture medium to the upper compartment of 3-µm-pore or 8-µm-pore polycarbonate membrane inserts (6.5 mm Transwell Permeable Supports cat. No. 3415 and 3422, Corning) coated with laminin at 2.9 x $10^4$ cells/well. Cells were allowed to transmigrate toward the lower compartment for 6 hours in the presence or absence of indicated drugs. Transmigration efficiency was calculated as number of cells at the lower compartment divided by the number of cells added to the upper compartment of a transwell.

Microfabrication-based devices

Micofluidic devices were fabricated in Polydimethylsiloxane (PDMS) using soft-lithographic procedures. A primary silicon mold with etched features (5 and 35 µm respectively for channels and expansion chambers) was prepared using 2 steps Reactive Ion Etching and aligned photolithography for patterning. PDMS, Sylgard 184 from Dow Corning, was prepared mixing the base resin and the reticulation agent in a 10:1 ratio. After vacuum degasing for about 30 min to remove all trapped air, the PDMS was poured on the silicon mold and cured for 1 hr at 70 °C on a hot plate. This PDMS first replica was then coated with a fluoro-silane (Trichloro-perfluoro silane, Sigma Aldrich 448931-10G) as an anti-sticking layer and used to produce the final devices in PDMS by cast molding: Sylgard 184 was ixed in a 10:1 ratio with reticulation agent and degassed, then poured on the PDMS mold an thermally cured for 1 hr at 70°C. The cured PDMS was then peeled-off the PDMS mold, and devices were diced by cutting using a razor blade. In- and Out-let point were punched through the PDMS with a 3 mm diameter circular puncher (Ted Pella, 15111-30 Biopunch), and finally each device was sealed to a glass coverslip using plasma activation of PDMS and glass surfaces to promote the chemical bonding. More detailed protocol is described in Supplementary Text.

Live-cell imaging

2D and 3D Matrigel cultures and dissociated CGCs on micropatterned substrates coated with laminin were observed with a spinning-disk confocal microscope CV1000 (Yokogawa) through a 20x objective (N.A. 0.75) or a 100x oil-immersion objective (N.A. 1.3), or with Dragonfly High Speed Confocal Microscopy System (Andor) through a 100x oil-immersion objective (N.A. 1.4), at 37°C with 5% $CO_2$ flow. For $Ca^{2+}$ imaging, the CGCs transfected with GCaMP6s were observed with an epifluorescent inverted microscope IX83 (Olympus) through a 100x oil-immersion objective (N.A. 1.3).

Westernblotting

Dissociated CGCs were plated in 60-mm cultured plates (Iwaki) coated with poly-D-lysine and laminin and maintained for 12 h in culture medium at 37°C/5% $CO_2$. LY333531 was added to cultures for 1 h at a final concentration of 10 µM before treatment with Yoda1. Yoda1 (Sigma) was added to cultures for 5 min at a final concentration of 100 µM. Equivalent volumes of solvent were added to control cultures. Cultures were washed with cold PBS and lysed in RIPA buffer (50 mmol/ L Tris-HCl (pH 7.6), 150 mmol/L NaCl, 1% NP40, 0.5% Sodium Deoxycholate, 0.1% SDS) with Protease inhibitor cocktail (EDTA free) (Nacalai) and Phosphatase inhibitor cocktail (Sigma). Lysates were incubated at 4 °C for 15 min and cleared at 15,000 × *g* for 15 min at 4 °C. Total protein was normalized using Protein assay BCA kit (Nacalai). Equal amounts of lysates were separated by SDS-PAGE in 4-20% Mini-PROTEAN TGX gel and transferred onto PVDF membrane (Millipore). Membranes were blocked for 60 min at room temperature in PBS-Tween with 5% skimmed powder milk. Membranes were incubated with primary antibodies in blocking buffer overnight at 4ºC at the following concentrations: anti-Ezrin (host, 1: 1/1000; Abcam), anti-phospho T567 ezrin (host, 1: 1/300; Abcam) and anti-GAPDH (host, 1: 1/2000; Abcam). Membranes were washed and incubated for 30 min in blocking buffer containing HRP-conjugated anti- rabbit or mouse secondary antibodies (1:10000; Biorad). Signal was detected with ECL Prime (G.E. Healthcare) and imaged on a ChemiDoc XRS+ System (Biorad).

BrdU pulse-labeling

5-bromo-2′-deoxyuridine (BrdU, Invitrogen) was dissolved in PBS. Wildtype and PIEZO1 conditional knockout mice at P9 were injected i.p. with BrdU (15 mg/kg) for labeling. Immunofluorescence and BrdU detection were done at P13.

Immunofluorescence

Mice were deeply anesthetized by isoflurane, and were transcardially perfused with phosphate buffer (PB) followed by 4% paraformaldehyde (PFA) in PB. Brains were removed and postfixed overnight in 4% PFA at 4°C. Brains were then washed of PFA with three changes of 1X phosphate buffered saline (PBS), and 30% sucrose/PBS. Dehydrated brains were frozen by liquid nitrogen, then embedded into O. C. T. compound (Sakura Finetek Japan). Sagittal sections (20 µm thick) were made using a cryostat (Leica). Slices on Super frost slide glasses (Matsunami) were treated with PBS-Triton X-100 (0.5% v/v) for 10 min, and washed with PBS-Triton X-100 (0.1% v/v). Brain slices were then blocked by incubation with 2% skim milk in PBS-Triton X-100 (0.1% v/v) for 30 min at RT. Primary antibody labeling was performed in blocking buffer overnight at 4°C as follows: anti-Pax6 (rabbit, 1:1000, Wako), anti-BrdU (mouse, 1:100, Sigma-Aldrich). After thorough washing, slices were incubated with secondary antibodies overnight at 4°C , followed by 10 ug/mL DAPI for 10 min at RT. After washing, slices were mounted with Fluoromount (DBS).

RNAscope

Fixed brains were transfered to 30% Sucrose/PBS and embedded in OCT compound. Cryosections (20 µm) were subjected to *in situ* hybridization according to the manufacturer's protocol (Advanced Cell Diagnostics, RNAscope). Probes were Mm-Piezo1-01(Cat No: 500511 LOT: 19100A) and DapB for negative control (Cat No: 310043 LOT: 2005245).

Quantitative image analyses and statistical analyses

Images analyses were performed using Fiji (Image J 2.0) or Python 3. MRLC-2GFP and ezrin-GFP signals in xy planar images resolved in 1 µm steps over the entire depth of the migrating cells were processed for maximum intensity projections. For the measurement of the intensity of cytoplasmic MRLC-2GFP, the signal intensity along a one pixel wide, 15 µm line near the midline of the cell soma was scanned in the front and rear of the nucleus. The intensity values were normalized to the average of all intensity values of each cell. The maximum intensity value obtained using the original python code were plotted for comparative analysis of data from different samples. For the measurement of the intensity of ezrin-EGFP on the plasma membrane, the signal intensity on the somal plasma membrane was scanned. The intensity values for the four subdivided areas (Front, Middle front, Middle rear, Rear) were normalized to the average of all intensity values in each cell. For quantification of $[Ca^{2+}]_i$, the average intensity of GCaMP6s signal in the soma was quantified manually in each time frame. The values were normalized to the average intensity in the soma before Yoda1 treatment.

Statistical analysis

Data were analyzed using one-way ANOVA with post-hoc Dunnett's test or the unpaired t test by GraphPad Prism 9 or R software.

**Supplementary Text**

1. Model of Neuronal Migration in Confined Space

2. Detailed Protocol for Microfabrication-based Devices

### 1. Model of neuronal migration in confined space

1-1. Simulation method

Computer simulation of a deformable spherical object that passes through a narrow tunnel was conducted based on the method of fluid-structure interaction simulation proposed in our previous studies (*5*, *6*). A spherical object was modeled as a slightly compressible object covered with a hyperelastic membrane. For simplicity, the intracellular structures such as the nucleus were not considered. The mechanical behavior of the two-dimensional membrane was assumed to be governed by Skalak's constitutive law (*7*), in which the strain energy function, $W$, is described using the first and second strain invariants of the right Cauchy–Green tensor, $I_1$ and $I_2$, respectively, as the following:

$$W = \frac{1}{4} G_\mathrm{s} \left( I_1^2 + 2I_1 - 2I_2 + CI_2^2 \right), \quad \text{(S1)}$$

where $G_\mathrm{s}$ is the surface shear elastic modulus, and $C$ is the area incompressibility coefficient. By using these two material properties, the surface Poisson's ratio, $\nu_\mathrm{s}$, and Young's modulus, $E$, can be expressed using the membrane thickness, $h$, as $\nu_\mathrm{s} = C/(C+1)$ and $E = 2G_\mathrm{s}(1+\nu_\mathrm{s})/h$, respectively. The contraction of the membrane, which mimics actomyosin contraction at the cell membrane, was modeled by introducing the active second Piola-Kirchhoff stress tensor, $\mathbf{S}_\mathrm{act}$, as

$$\mathbf{S}_\mathrm{act} = A\boldsymbol{m}_0 \otimes \boldsymbol{m}_0, \quad \text{(S2)}$$

where, $A$ is the magnitude of contraction and $\boldsymbol{m}_0$ is the unit vector representing the direction of contraction in the reference state; thereby this model expresses unidirectional contraction. The membrane of the spherical object was discretized into triangular finite elements. The restoring force, $\boldsymbol{q}_\text{in-plane}$, acting on the membrane node $\boldsymbol{x}_\mathrm{m}$ due to the membrane deformation was computed based on a finite element procedure.

To consider a volume constraint of the spherical object and a bending stiffness of the membrane, we introduced the Helmholtz free energies, $F_\mathrm{vol}$ and $F_\mathrm{bend}$, respectively (*8*). The free energy for the volume constraint, $F_\mathrm{vol}$, was proposed using the object volume in the current state, $V$, as

$$F_\mathrm{vol} = \frac{1}{2} k_\mathrm{vol} \left( \frac{V}{V_0} - 1 \right)^2, \quad \text{(S3)}$$

where $k_\mathrm{vol}$ is the bulk modulus and $V_0$ is the object volume in the reference state, i.e., the volume of the undeformed sphere. Since this is an artificial free energy to model the effect of slightly compressible internal fluid, the exact value of $k_\mathrm{vol}$ is unimportant. Similarly, the bending free energy, $F_\mathrm{bend}$, was expressed using the angle between two adjacent triangular elements ($\alpha$, $\beta$) in the current state, $\theta_{ab}$, as

$$F_\mathrm{bend} = \sum_{\text{adjacent}\,\alpha,\beta\,\text{pair}} k_\mathrm{bend} \left[ 1 - \cos\left( \theta_{\alpha\beta} - \theta_0 \right) \right], \quad \text{(S4)}$$

where, $k_{\text{bend}}$ is the bending modulus and $\theta_0$ is the angle in the reference state, which was set to 0 in this study. These free energies can produce the corresponding restoring forces, $\boldsymbol{q}_{\text{vol}}$ and $\boldsymbol{q}_{\text{bend}}$, on the membrane node $\boldsymbol{x}_{\text{m}}$ as the following:

$$\boldsymbol{q}_{\text{vol}}\left(\boldsymbol{x}_{\text{m}}\right) = -\frac{\partial F_{\text{vol}}}{\partial \boldsymbol{x}_{\text{m}}}, \tag{S5}$$

$$\boldsymbol{q}_{\text{bend}}\left(\boldsymbol{x}_{\text{m}}\right) = -\frac{\partial F_{\text{bend}}}{\partial \boldsymbol{x}_{\text{m}}}. \tag{S6}$$

Additionally, by considering externally applied force, $\boldsymbol{q}_{\text{ext}}$, on the membrane node $\boldsymbol{x}_{\text{m}}$, the total force, $\boldsymbol{q}$, acting on $\boldsymbol{x}_{\text{m}}$ is given by

$$\boldsymbol{q}\left(\boldsymbol{x}_{\text{m}}\right) = \boldsymbol{q}_{\text{in-plane}}\left(\boldsymbol{x}_{\text{m}}\right) + \boldsymbol{q}_{\text{vol}}\left(\boldsymbol{x}_{\text{m}}\right) + \boldsymbol{q}_{\text{bend}}\left(\boldsymbol{x}_{\text{m}}\right) + \boldsymbol{q}_{\text{ext}}\left(\boldsymbol{x}_{\text{m}}\right). \tag{S7}$$

The proposed spherical object model was placed in a channel filled with Newtonian fluid. The fluid dynamics was numerically analyzed using the lattice Boltzmann method (*9*). Furthermore, the membrane deformation of the spherical object and the surrounding fluid dynamics were coupled using the immersed boundary method (*10*). In this method, the membrane force $\boldsymbol{q}$ (Eq. S7) is distributed to the neighboring fluid to drive the fluid flow, and conversely, the fluid velocity determines the neighboring membrane velocity for the advection.

1-2. Simulation model

A three-dimensional model of two cylindrical channels connected by a narrow cylindrical tunnel was constructed (Fig. 1G). Each channel has a diameter of 12 μm and a length of 12 μm, and the connecting tunnel has a diameter of 4 μm and a length of 4 μm. The fluid-filled space in the channels and tunnel was discretized using a three-dimensional regular lattice with an interval of 0.4 μm. The spherical object model proposed in the previous section was placed in the center of one cylindrical channel. The membrane was discretized using two-dimensional triangular finite elements with an approximately 0.2 μm edge size, which was set to be smaller than the lattice interval in the fluid-filled space to avoid leaks. To move the spherical object forward, a tensile force of 60 pN was applied uniformly on the front membrane, which is equivalent to a solid angle of 0.1 π steradian (green region shown in Fig. 1G), along the longitudinal direction of the channels.

After the deformed spherical object ceased to move in the tunnel, contraction of the rear hemispherical membrane (red region shown in Fig. 1G) with the magnitude of $A = 5.0 \times 10^{-4}$ (Pa·m) was imposed (see Eq. S2), where the direction of contraction $\boldsymbol{m}_0$ was randomly set for each triangular element within its tangential plane to represent in-plane isotropic contraction of the membrane. To verify the effectiveness of the membrane contraction to pass through the narrow tunnel, the simulation without contraction was also performed.

As boundary conditions, the no-slip condition was applied at the wall of the channels and tunnel, and the periodic boundary condition for the fluid velocity was applied at both ends of the channels. No frictional force was assumed between the wall of the tunnel and the membrane of the spherical object. The material properties of the spherical object used in the present simulation are listed in Table S1 (*11*, *12*). By using these material properties, the surface shear elastic modulus, $G_s$, and the surface Poisson's ratio, $\nu_s$, of the membrane can be derived as $G_s = 5.2 \times 10^{-7}$ (Pa·m) and $\nu_s = 0.91$, respectively.

**2. Detailed Protocol for Microfabricated-based Devices**

2-1. Materials

Silicon wafers:
4” Single side polished silicon wafer, with thermal oxide (300 nm thick) on both sides

Photoresist:
AZ5214E (Merck Performance Materials Pte Ltd, Singapore)

Developer:
AZ 400K (Merck Performance Materials Pte Ltd, Singapore)

Poly(dimethylsiloxane) (PDMS):
Sylgard 184 Silicone Elastomer Kit

Perfluorosilane:
Trichloro(1H,1H,2H,2H-perfluorooctyl)silane
(Merck, CAS Number 78560-45-9, Product number 448931)

DI Water:
Milli-Q (Merck Millipore)

Isopropyl-alchool
EL grade (CAS Number: 67-63-0, Aik Moh Singapore)

Acetone
EL grade (CAS Number: 67-64-1, Aik Moh Singapore)

Process Gases:
$O_2$, $CF_4$, $C_4F_8$, $SF_6$ (Air Liquide Singapore PTE ltd, 99.9995% purified)

2-2. Methods

I. First lithography step:
We used 4”, single-side polished (100) prime Si wafers with 300 nm thick thermally grown $SiO_2$ wafers for the fabrication of the master mold.

A 1.5 µm thick layer of positive tone photo-resist (AZ5214E) is deposited on the polished side of the wafer by spin-coating at 2500 rpm for 45 s. After spin-coating, a pre-bake is applied to remove the excess of solvent (90 s at 110 °C on a Hot Plate). The photo-resist is then exposed to UV light through an optical mask with the pattern of the final device (about 100 mJ/cm$^2$ energy dose, UV photons at 365 nm generated by a 500 W Xe-Hg arc lamp) and developed to reveal the exposed image by immersion in AZ 400K developer diluted 1:4 in DI water for 40 s. After rinsing with clean DI water and drying with blowing $N_2$ gas, the wafer is ready for the next step.

II. Oxide layer dry etching:

The 300 nm thick oxide layer is etched in a Reactive Ion Etching chamber using the patterned photo-resist as a mask. We used a Samco RIE-10NR tool for the silicon oxide etching with the following process parameters: 150 W power (source at 13.56 MHz), gas mixture of $CF_4$ (40 sccm) and $O_2$ (4 sccm) at a chamber pressure of 15 Pa; this process has an etching rate of approx. 100 nm/min in $SiO_2$, thus we applied the process for 3 min 15 s where the small excess of time is to make sure the oxide layer is fully removed from the exposed areas.

III. Residual photo-resist removal:

The photo-resist remainng after the oxide etching is completely removed and the wafer is cleaned in two step; first the wafer is kept in a bath of Acetone for 5 min with gentle agitation, then the wafer is moved in two a bath of Isopropyl alcohol and washed there for 30 s, finally the wafer is removed from the bath and rinsed with $N_2$ blow gun. The second and final cleaning step is an $O_2$ plasma process which we carried in a Diener Pico plasma tool, with 20 sccm $O_2$ flow, the chamber is kept at 2 mbar pressure and a power of 90 W is applied to the gas to generate the plasma for 2 min.

IV. Second lithography:

On the cleaned wafer, AZ5214E photo-resist is coated in the same way as in step 1, then lithography is done with a different optical mask with the second lay-out. The lithographic process is exaclty the same as in step 1, the patterned photo-resist will leave exposed only selected portions of the previous pattern etched in the silicon oxide layer.

V. First deep silicon etching:

In this step, the wafer as prepared after step 3 is plaved in the reactor of an Inductively Coupled Plasma machine (ICP) for a first step of deep silicon etching. Silicon etching in an ICP is capale of producing vertical profiles with etching depth up to several tens of µm. We used a BOSH-like process, which works with alternating a coating step with an etching one, resulting in vertical pattern transfert. The processes for the two steps are, respectively:

a. Coating step: $C_4F_8$ (140 sccm) is fluxed in the vacuum chamber kept at a pressure of 6 Pa and 500 W of power (source at 13.56 MHz) are applied to generate the plasma for a duration of 6 s. The plasma in this step produces a thin teflon-like comformal coating on the wafer
b. Etching step: $SF_6$ ( 130 sccm) and $O_2$ (13 sccm) gas mixture is fluxed in the vacuum reactor chamber kept at 8 Pa of pressure and 500 W of power (source at 13.56 MHz) are applied to generate the plasma for a duration of 8 s. The plasma in this step produces reactive species which remove the thin protective coating generated in the step a mor eefficiently on the horizontal surfaces and then chemically remove Si where exposed

This etching process in our Sentech 500Si ICP reactor produces an etching rate in Si of approximately 250 nm/cycle (1 cycle: 1 coating step followed by 1 etching step). In this first silicon etching step we reached a final etching depth of 30 µm (120 total cycles)

VI. Photo-resist removal:

After the first silicon etching, the wafer is removed from the ICP and the residual photo-resist mask is cleaned similarly to step 3. At this stage, we confirmed the ethced depth in the silicon with a Bruker Stylus profiler.

VII. Second and final silicon etching:

The wafer affter step 5 is placed back in the same ICP reactor, and the same process is applied to etech extra 5 µm in the Si (20 cycles). The previous feature will results finally in a 35 µm deep cavityes, while the feature exposed for the frist time will have a final etch depth of 5 µm.

VIII. Silanization:

The wafer after step 6 is patterned with fetures etched down to a final depth of 35 and 5 µm. This wafer is now considered as the Master Mold for replica molding with PDMS. In this step the mold is treated with an anti-sticking coating by vapour-deposition of a molecular layer of Trichloro(1H,1H,2H,2H-perfluorooctyl)silane. First, the surface of the mold is acitvated with a 30 s $O_2$ plasma (Pico Diener plasma tool, $O_2$ 20 sccm with the chamber at 2 mbar presure, 90 W RF power at 13.56 MHz) then immediately placed in a plastic vacuum jar together with a small drop of perfluorosilane (approx. 20 µL) and kept in vacuum (1 mbar or less) for at least 1 hr. After at least 1 hr, the wafer can be retrieved from the vacuum jar and it is ready to be used for PDMS casting multiple times.

IX. PDMS replica:

Sylgard 184 PDMS is prepared by mixing the silicone elastomer with his curing agent in a 10:1 ratio (in weight, prepare about 25 g total). After mixing, degas in a vacuum jar (approx 1 mbar) for about 30 min, or until no more visible air bubbles are visible.

Place the silanized master mold in a petri dish and slowly pour the degassed PDMS until the master if completely covered. Place the petri dish containing the master mold covered with PDMS in the vacuum jar and degas at 1 mbar or lower for 5-10 min to ensure no air is trapped at the surface of the mold.

Place the petri dish with the mold and the PDMS on a hot plate at 70 °C or in an oven at 70 °C an allow for the PDMS to thermally cure for about 1.5 hr. After thermal curing is done, with a blade carefully cut around the edge of the mold then slowly peel-off the cutted section of cured PDMS from the mold (which is left in the petri dish). Store the PDMS cut in a second petri dish; the master mold can be kept in its petri dish sourrounded by cured PDMS and protected with the petri dish lid. The silicon master mold can be re-used for multiple steps of replica molding.

X. PDMS working mold preparation:

The PDMS replica of the master mold prepared in step 8 is now patterned with protrusion that replicate exactly, the features in the master mold (with inverted polarity). This PDMS replicas can be used as a Working Mold to produce by a second step of PDMS replica molding the final devices. Prior to PDMS casting on the Working Mold, it needs to be coated with the same anti-sticking layer as in step 7. The process is identical, except the $O_2$ plasma is carried on at lower poew (30 W).

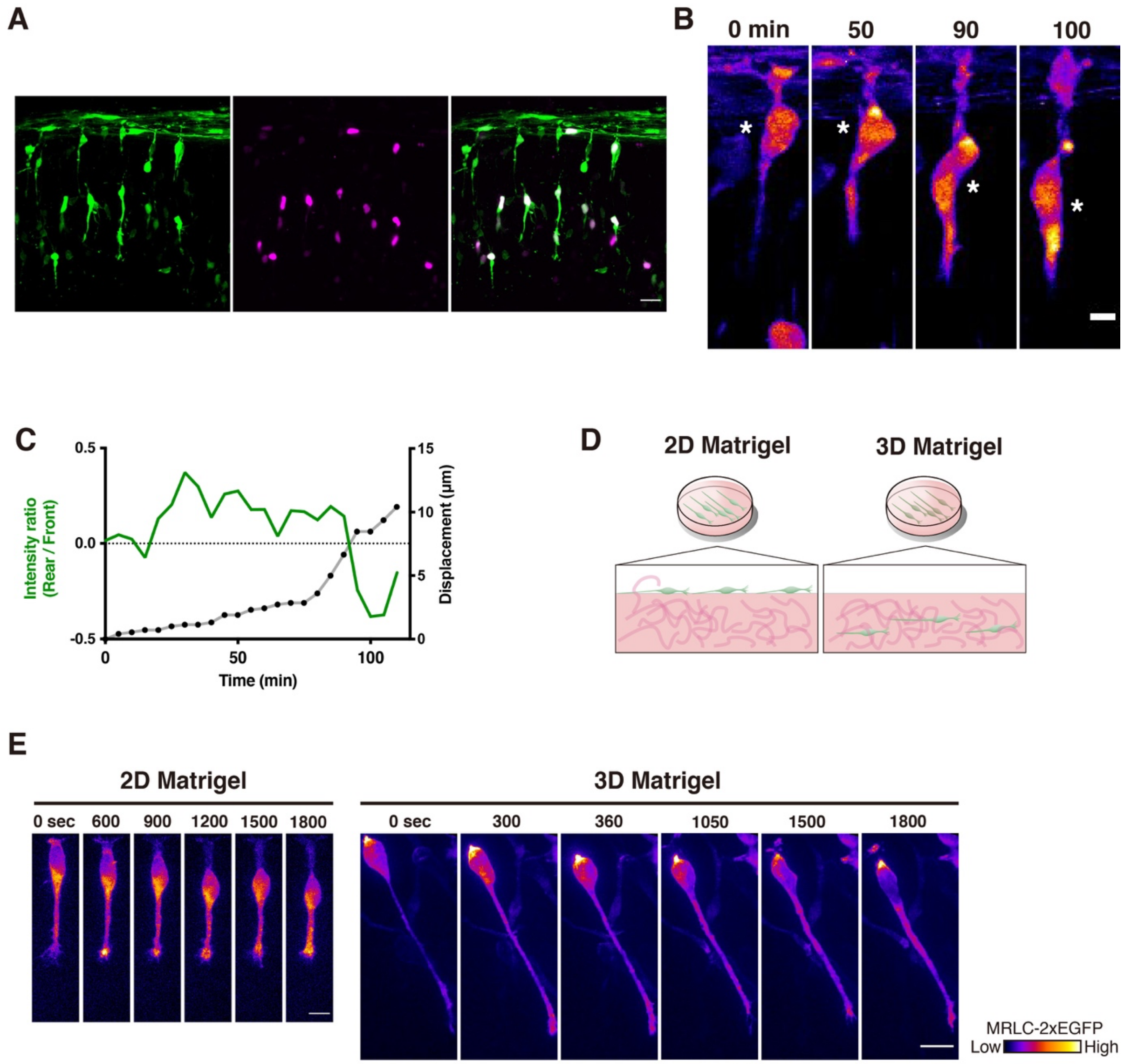


Fig. S1, Nakazawa et al.,

**Fig. S1. Dynamic changes in myosin localization in CGNs and cortical interneurons in 2D and 3D cultures**

(**A**) Snapshot image of CGNs electroporated with MRLC-2GFP (green) and mScarlet-NLS (magenta) in the cerebellar slice. (**B**) Image sequence of another CGN migrating in the cerebellar tissue from a P10 mouse. (**C**) Position of the soma (approximate oval center indicated by asterisks in B) and relative (rear/front) distribution of myosin (MRLC-GFP) in the neuron shown in S1B plotted against time. (**D**) 2D and 3D CGN cultures. (**E**) Image sequence of myosin localization in migrating cortical interneurons in 2D and 3D Matrigel culture. Scale bars, (A), 20 µm; (B), 5 µm; (E), 10 µm.

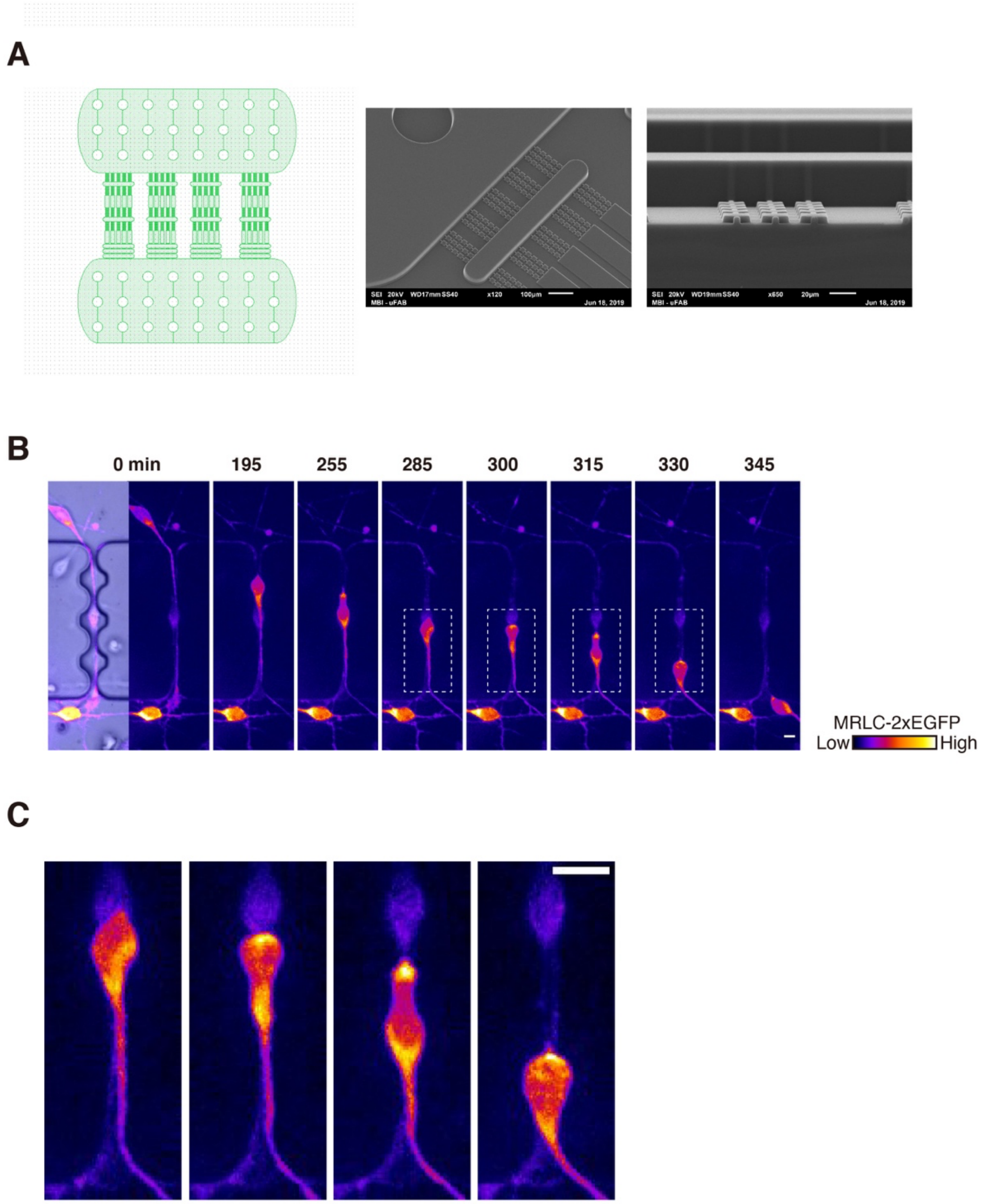


Fig. S2, Nakazawa et al.,

**Fig. S2. Actomyosin dynamics in CGNs migrating in confined spaces in microfluidic channels**

(**A**) Scheme of microfluidic devices used to observe CGN migration in repetitive narrow channels. (**B**) A bright field image (left) and image sequence of a CGN labeled with MRLC-2GFP migrating in the microchannels (right 7 images). (**C**) Magnified view of the CGN in (**B**) at indicated time points. Scale bars in (B) and (C), 5 µm.

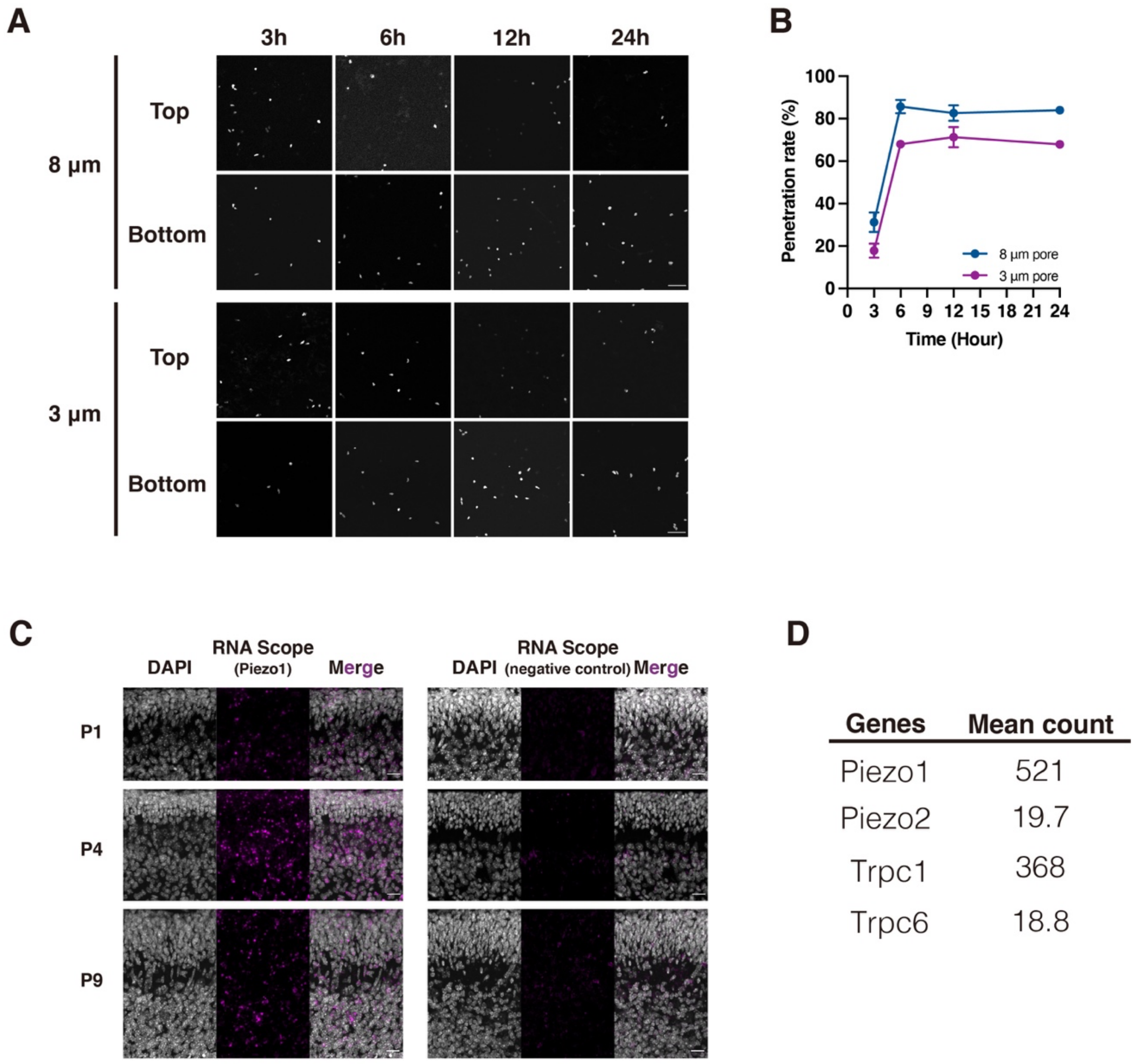


Fig. S3, Nakazawa et al.,

**Fig. S3. Transwell migration assay of CGNs**

(**A**) CGNs on the top and bottom of the porous membranes after transwell migration. CGNs were inoculated on top of the 3-µm or 8-µm pore membrane and fixed at indicated time. (**B**) The percentages of CGNs that have completed transwell migration at indicated time after inoculation. Samples were collected from three independent experiments. (**C**) RNAscope in situ hybridization for PIEZO1 mRNA in developing cerebellum at P1, P4, and P9. EGL; external granule layer, ML; molecular layer, IGL; internal granule layer. Scale bars, 50 µm. (**D**) Expression of PIEZO1 and PIEZO2 in CGNs from P6 mice as revealed by RNA-seq. See also Data S1.

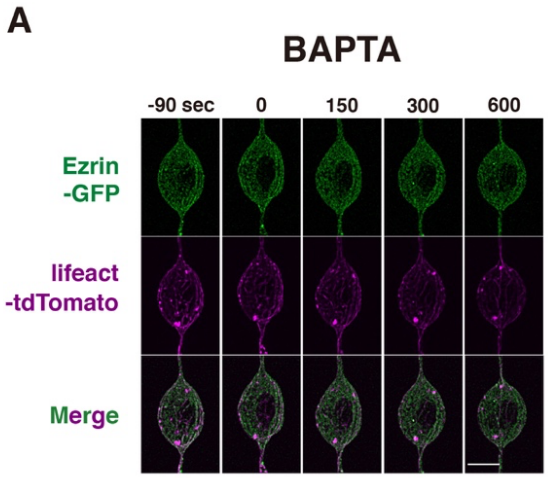


Fig. S4, Nakazawa et al.,

**Fig. S4. Piezo1-induced translocation of ezrin and actomyosin is mediated by $Ca^{2+}$influx and PKC**

Time-lapse sequence of ezrin-GFP and lifeact-tdTomato in CGNs upon Yoda1 treatment in the presence of BAPTA (**A**) or LY333531 (**B**). Scale bars, 5 µm.

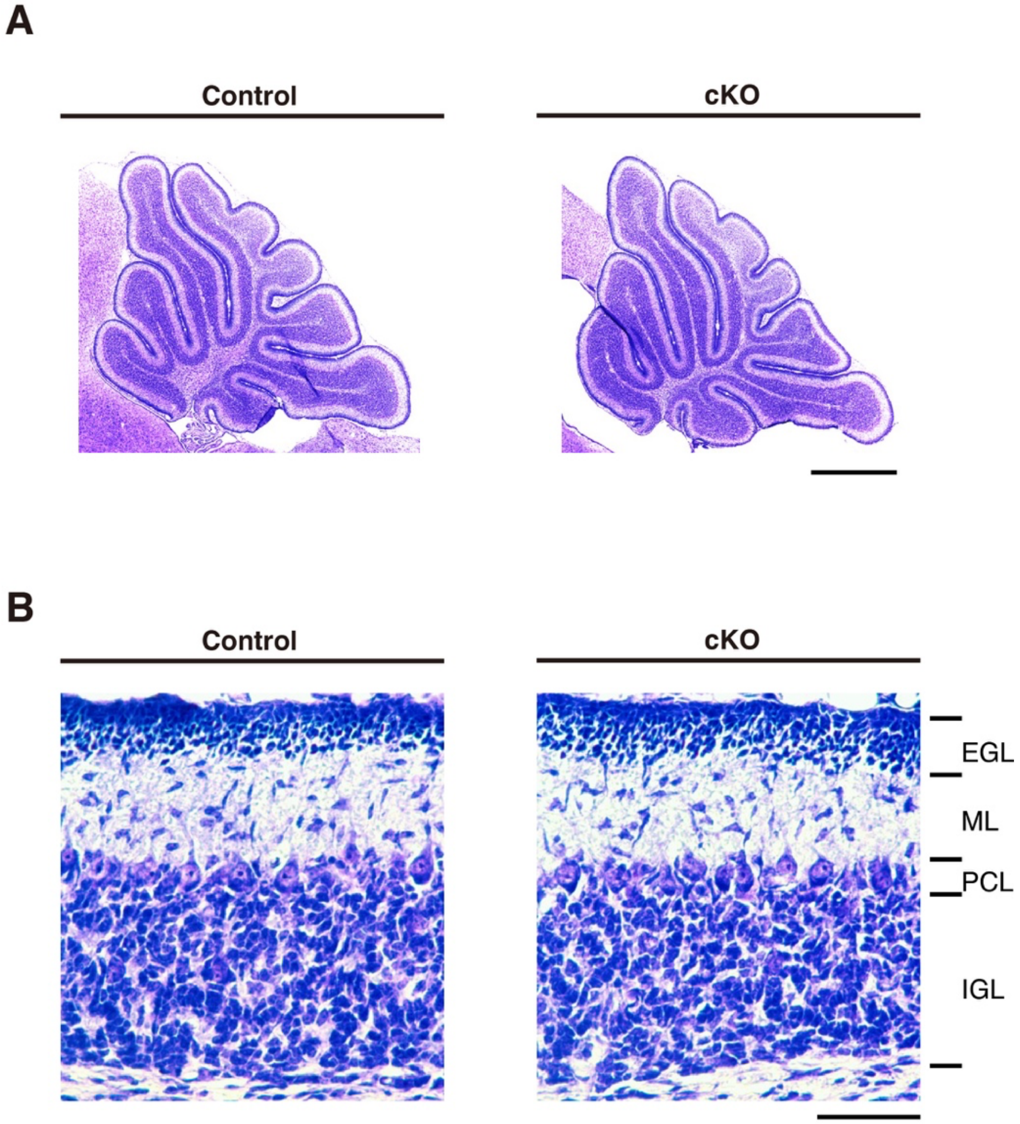


Fig. S5, Nakazawa et al.,

**Fig. S5. Apparent normal formation of the cerebellar cortex in PIEZO1 cKO mice**
(**A**) Sagittal cerebellar sections of control (Piezo1 $^{flox/flox}$) and PIEZO1 cKO (NeuroD1-Cre; Piezo1$^{flox/flox}$) mice at P9. (B) Magnified views of the cerebellar cortices. Scale bars, A, 500 μm; B, 50 μm.

**Table S1. Material properties of the spherical object for the fluid–structure interaction simulation**.

| Symbol (unit) | Description | Value |
|---|---|---|
| $E$ (Pa) | Young's modulus of membrane | 200 (*8*) |
| $C$ | Area incompressibility coefficient | 10 |
| $h$ (nm) | Thickness of membrane | 10 |
| $k_{vol}$ (J) | Bulk modulus of spherical object | $1.0 \times 10^{-13}$ |
| $k_{bend}$ (J) | Bending modulus of membrane | $5.0 \times 10^{-20}$ (*11*) |

### Reference for Supplementary Materials


1. S. M. Cahalan, V. Lukacs, S. S. Ranade, S. Chien, M. Bandell, A. Patapoutian, Piezo1 links mechanical forces to red blood cell volume. *Elife*. **4** (2015), doi:10.7554/eLife.07370.

2. M. Sawada, M. Matsumoto, K. Narita, N. Kumamoto, S. Ugawa, S. Takeda, K. Sawamoto, In vitro Time-lapse Imaging of Primary Cilium in Migrating Neuroblasts. *Bio Protoc*. **10**, e3823 (2020).

3. Y. K. Wu, H. Umeshima, J. Kurisu, M. Kengaku, Nesprins and opposing microtubule motors generate a point force that drives directional nuclear motion in migrating neurons. *Development*. **145** (2018), doi:10.1242/dev.158782.

4. H. Umeshima, T. Hirano, M. Kengaku, Microtubule-based nuclear movement occurs independently of centrosome positioning in migrating neurons. *Proc. Natl. Acad. Sci. U. S. A.* **104**, 16182–16187 (2007).

5. Y. Kameo, M. Ozasa, T. Adachi, Computational framework for analyzing flow-induced strain on osteocyte as modulated by microenvironment. *J. Mech. Behav. Biomed. Mater.* **126**, 105027 (2022).

6. Y. Yokoyama, Y. Kameo, H. Kamioka, T. Adachi, High-resolution image-based simulation reveals membrane strain concentration on osteocyte processes caused by tethering elements. *Biomech. Model. Mechanobiol.* **20**, 2353–2360 (2021).

7. R. Skalak, A. Tozeren, R. P. Zarda, S. Chien, Strain energy function of red blood cell membranes. *Biophys. J.* **13**, 245–264 (1973).

8. J. Li, M. Dao, C. T. Lim, S. Suresh, Spectrin-level modeling of the cytoskeleton and optical tweezers stretching of the erythrocyte. *Biophys. J.* **88**, 3707–3719 (2005).

9. S. Chen, G. D. Doolen, Lattice Boltzmann method for fluid flows. *Annu. Rev. Fluid Mech.* **30**, 329–364 (1998).

10. C. S. Peskin, The immersed boundary method. *Acta Numer.* **11**, 479–517 (2002).

11. M. Puig-de-Morales-Marinkovic, K. T. Turner, J. P. Butler, J. J. Fredberg, S. Suresh, Viscoelasticity of the human red blood cell. *Am. J. Physiol. Cell Physiol.* **293**, C597-605 (2007).

12. E. Spedden, J. D. White, E. N. Naumova, D. L. Kaplan, C. Staii, Elasticity maps of living neurons measured by combined fluorescence and atomic force microscopy. *Biophys. J.* **103**, 868–877 (2012).

**Data S1.**

RNA-seq analysis of P6 mouse cerebellum.
https://drive.google.com/file/d/1dVX1I5fzlq2yCVpMELAR5c1vfkaCappV/view?usp=sharing
